# A statistical approach to genome size evolution: Observations and explanations

**DOI:** 10.1101/031963

**Authors:** Dirson Jian Li

## Abstract

Genome size evolution is a fundamental problem in molecular evolution. Statistical analysis of genome sizes brings new insight into the evolution of genome size. Although the variation of genome sizes is complicated, it is indicated that the genome size evolution can be explained more clearly at taxon level than at species level. I find that the genome size distribution for species in a taxon fits log-normal distribution. And I find a relationship between the phylogeny of life and the statistical features of genome size distributions among taxa. I observed different statistical features of genome size distributions between animal taxa and plant taxa. A log-normal stochastic process model is developed to simulate the genome size evolution. The simulation results on the log-normal distributions of genome sizes and their statistical features agree with the observations.

## Introduction

Genome size evaluates the total amount of DNA contained within the genome of a species. Genome sizes vary greatly among species. The genome size evolution is a fundamental problem in molecular evolution, which concerns the macroevolution of life (Gregory 2004). Several models were proposed to explain the genome size evolution from different perspectives (Petrov 2002, Karev et al. 2003, Lygeros et al. 2008). But the mechanism for genome size evolution is not clear, and the distributional pattern of genome sizes remains unknown (Gregory 2005). There is more non-coding DNA in eukaryotes than in prokaryotes, so the genome size variation is more complicated for eukaryotes than for prokaryotes. It seems that genome size does not correlate with organism complexity, some simple organisms have more DNA than complex ones due to the varying amount of non-coding DNA. However, it is still hard to explain the origin and evolution of non-coding DNA (Castillo-Davis 2005, Bird et al 2006, Taft et al. 2007). The term C-value enigma (Gregory 2001, Gregory 2002) was coined to refer these problems on genome size evolution, especially on non-coding DNA. Understanding the C-value enigma will shed light on the macroevolution of life at molecular level.

Genome size evolution is a complex process, which is driven by either the evolutionary factors or the ecological factors. Both small- and large-scale duplication events have played significant roles in genome size evolution (Gregory 2005). Gene family expansion is an ongoing phenomenon, and large gene families are a feature of metazoan genomes. Polyploidy plays an especially important role in plant genome evolution. Polyploidy is also a widespread phenomenon in the animal kingdom. Transposable elements often contribute significantly to genome size changes. The mechanisms for amplification and spread of transposable elements are transposition, horizontal transfer, and sexual reproduction. Loss of transposable elements can also occur. Although the overall trend of genome size evolution has been in the direction of increase, the change of genome size is not a uni-directional process. In many cases, the genome size will shrink back very quickly after whole genome duplication due to selective constrains. There are also some groups of animals whose genome underwent extensive gene loss. Through the long evolutionary history, genome size evolution can be taken as a stochastic process due to the above multi-factorial drivers and their complexity. The distribution of genome sizes of the closely-related species in a taxon may fit certain statistical distribution. The statistical features of genome sizes reveal insight into their evolutionary history.

The main purpose of this paper is to explore the mechanism for genome size evolution. I provide statistical analysis of genome sizes at taxon level rather than at species level. Although the variation of genome sizes seems complicated at species level, I observe some statistical features of genome sizes of species among taxa. I find that the genome sizes of species in a taxon generally fit log-normal distribution. Genome size layout is defined for a certain taxon based on the relationship between the genome size variation ranges and the standard deviations of logarithmic genome sizes for all the included taxa of lower rank (Fig 1C, 1D). The genome size layout for a taxon reveals its evolutionary history. It is observed that the smaller the standard deviation of logarithmic genome sizes is, the smaller/greater the logarithmic mean of genome sizes for Dicotyledoneae/Monocotyledoneae taxa is (Fig 1D). So the genome size layout for angiosperm is V-shaped (Fig 1D). Based on biological data, I observe Λ-shaped genome size layout for animal and V-shaped genome size layout for plant respectively. It is indicated that the genome size evolution follows a log-normal stochastic process. A log-normal stochastic process model is developed to simulate the statistical features of genome sizes of species among taxa. The log-normal distribution of genome sizes and the different types of genome size layouts are simulated by the model (Fig 2), by which the relationship between the phylogeny of life and the statistical features of genome sizes are explained. The statistical features of genome sizes for animal and plant in simulations (Fig 3) agree with the observations (Fig 1).

**Figure 1:**
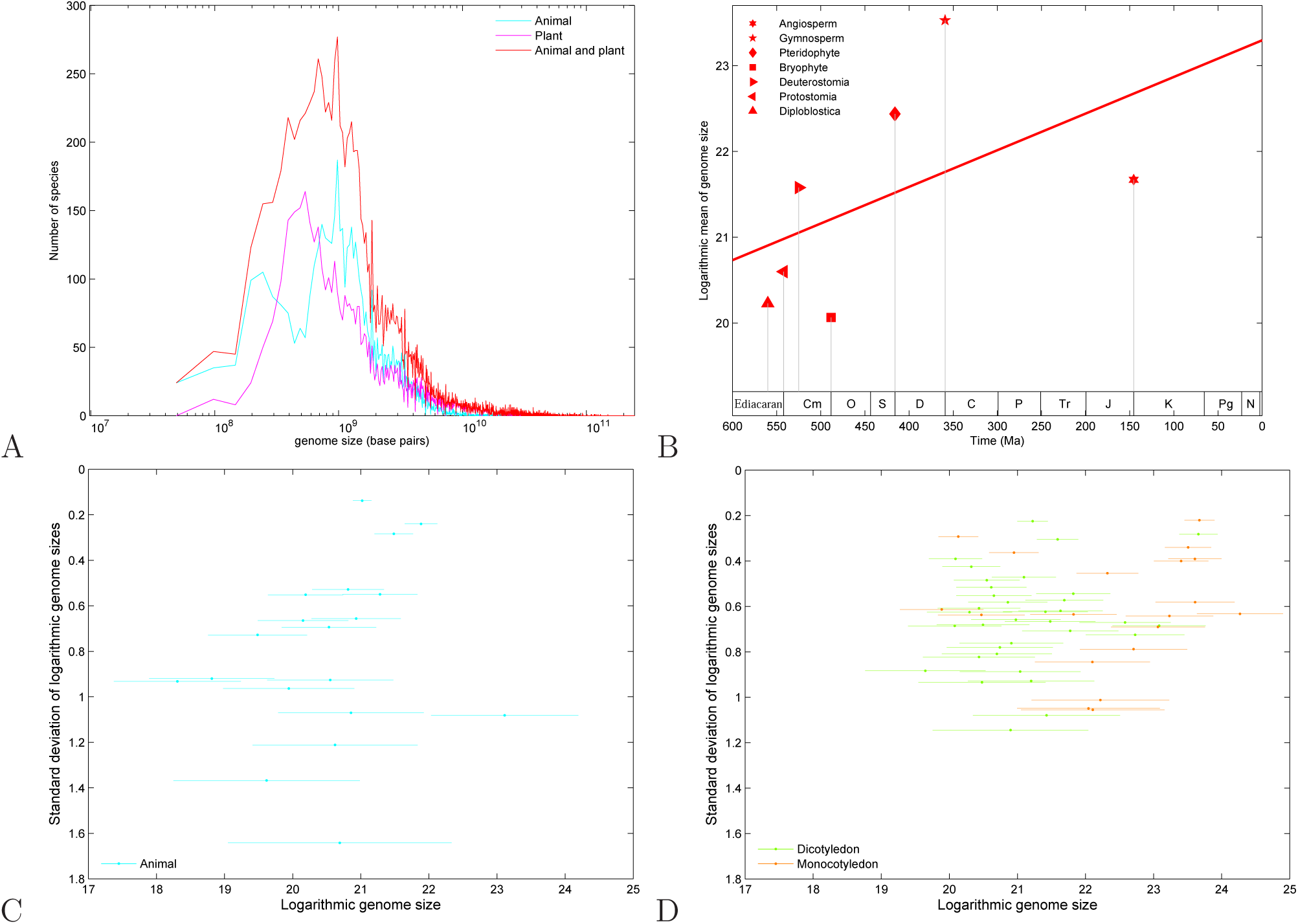
Statistical features of genome sizes among taxa in observation, based on genome sizes of contemporary species. **A** Log-normal distributions of genome sizes for animal, plant, and both of them in observation. **B** Exponential growth trend in genome size evolution through the Phanerozoic eon, according to the roughly linear relationship between the origin time of the 7 taxa and their logarithmic means of genome sizes. **C** Genome size layout for animal, which is about Λ-shaped. **D** Genome size layout for angiosperm, which is V-shaped. Most Dicotyledon taxa are on the left, while most Monocotyledon taxa on the right.

**Figure 2:**
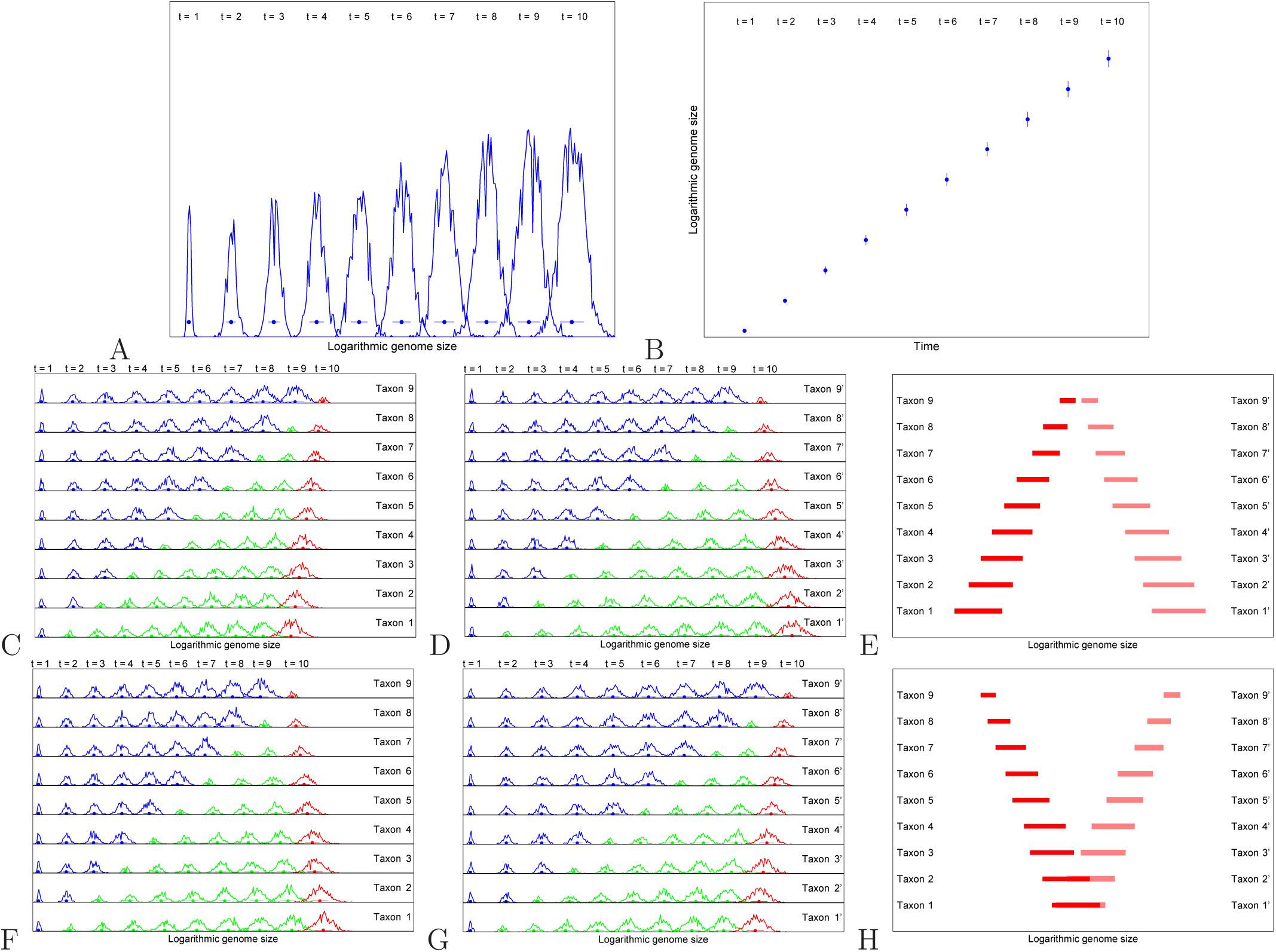
Simulation of the statistical features of genome sizes among taxa based on a log-normal stochastic process model. (A~B) Simulation of the genome size evolution for a taxon based on the log-normal stochastic process model. **A** Simulation of the log-normal distributions for a taxon at different stages *t* = 1,2,…, 10 in genome size evolution. **B** Simulation of the exponential growth trend in genome size evolution based on the linear relationship between time *t* and the logarithmic means of genome sizes simulated in Fig 2A, which agrees with the linear relationship between time *t* and the logarithmic means of genome sizes in observation (Fig 1B). (C ~ E) Simulation of the Λ-shaped genome size layout. **C** Simulation of the left part of the Λ-shaped genome size layout, where the probability for ancestor taxa is constant *P_a_* = 0.2, and the probability for descendant taxa is *P_d_* = 0.175, which corresponds to a relatively slow genome size growth rate for the descendant taxa. The ancestor taxa are in blue, while the descendant taxa are in green; especially the taxa at time *t* = 10 represents the modern or the present taxa, which are in red (the same colour convention below). **D** Simulation of the right part of the Λ-shaped genome size layout, where the probability for ancestor taxa is constant *P_a_* = 0.2, and the probability for descendant taxa is *P_d_* = 0.225, which corresponds to a relatively fast genome size growth rate for the descendant taxa. **E** Simulation of the Λ-shaped genome size layout based on the results at time *t* = 10 in Fig 2C and 2D. (F~H) Simulation of the V-shaped genome size layout. **F** Simulation of the left part of the V-shaped genome size layout, where the probability for descendant taxa is constant *P_d_* = 0.2, and the probability for ancestor taxa is *P_a_* = 0.175, which corresponds to a relatively slow genome size growth rate for the ancestor taxa. **G** Simulation of the right part of the V-shaped genome size layout, where the probability for descendant taxa is constant *P_d_* = 0.2, and the probability for ancestor taxa is *P_a_* = 0.225, which corresponds to a relatively fast genome size growth rate for the ancestor taxa. **H** Simulation of the V-shaped genome size layout based on the results at time *t* = 10 in Fig 2F and 2G.

## Materials and Methods

### Data resources

The genome sizes of animals are from the Animal Genome Size Database, and the genome sizes of plants are from the Plant DNA C-values Database (release 5.0, Dec. 2010) (Gregory et al. 2007). In this paper, we generally follow the taxonomy in the Animal Genome Size Database and the taxonomy in the Plant DNA C-values Database.

Based on the taxonomy in the Animal Genome Size Database and the Plant DNA C-values Database (Gregory et al. 2007), 7 taxa are taken as sample taxa for Eukarya; 19 taxa are taken as sample taxa for animal; 53 taxa are taken as sample taxa for angiosperm, where there are 34 Dicotyledoneae taxa and 19 Monocotyledoneae taxa. Concretely, the 7 Eukaryotic taxa are as follows: Diploblastica. Protostomia, Deuterostomia, Bryophyte, Pteridophyte, Gymnosperm, and Angiosperm, where there are 3 animal higher taxa and 4 plant higher taxa. The 19 animal taxa are as follows: Ctenophores, Sponges, Cnidarians, Anthropod, Molluscs, miscellaneous Invertebrates, Nematodes, Flatworms, Annelid, Myriapods, Rotifers, Tardigrades, Amphibian, Bird, Fish, Mammal, Reptile, Echinoderm, and Chordates. Among the 53 angiosperm taxa, the 34 Dicotyledoneae taxa are as follows: Lentibulariaceae, Cruciferae, Rutaceae, Oxalidaceae, Crassulaceae, Rosaceae, Boraginaceae, Labiatae, Vitaceae, Cucurbitaceae, Onagraceae, Leguminosae, Myrtaceae, Polygonaceae, Euphorbiaceae, Convolvulaceae, Chenopodiaceae, Plantaginaceae, Rubiaceae, Caryophyllaceae, Amaranthaceae, Malvaceae, Umbelliferae, Solanaceae, Papaveraceae, Compositae, Cactaceae, Passifloraceae, Orobanchaceae, Cistaceae, Asteraceae, Ranunculaceae, Loranthaceae, and Paeoniaceae, and the 19 Monocotyledoneae taxa are as follows: Cyperaceae, Juncaceae, Bromeliaceae, Zingiberaceae, Iridaceae, Orchidaceae, Araceae, Gramineae, Palmae, Asparagaceae, Agavaceae, Hyacinthaceae, Commelinaceae, Amaryllidaceae, Xanthorrhoeaceae, Asphodelaceae, Alliaceae, Liliaceae, and Aloaceae.

### Observation of log-normal distribution of genome sizes

Genome sizes vary greatly. The variation ranges of genome sizes of species in taxa often span across several orders of magnitude. Statistical analysis of genome sizes is applied to each of the above taxa in this work. Genome size distribution is defined as the distribution of genome sizes of species in a certain taxon. I observed that the genome size distribution is skewed: the left side of the peak is steep, while the right side of the peak is gentle. Obviously, the genome sizes of species in a taxa do not fit normal distribution. However, the logarithm of genome sizes of species in a taxa often fit normal distribution. So it is indicated that the genome sizes of spices in a certain taxon follow log-normal distribution. A log-normal distribution is a continuous statistical distribution of a random variable whose logarithm is normally distributed. The log-normal distribution is important in the description of some biological phenomena. For instance, sizes of organisms are often distributed log-normally. It can be shown that a growth process, whose relative growth rate is independent of size, will result in entity sizes with a log-normal distribution (Sutton 1997). In the case of genome size evolution, the growth rate of genome size is generally independent of genome size, so the genome sizes of spices in a certain taxon generally follow log-normal distribution.

I calculate the genome size distributions for the 7 Eukaryotic taxa, the 19 animal taxa, and the 53 angiosperm taxa, respectively. The genome size distributions for these taxa approximately fit lognormal distribution. I obtain the genome size distribution for animal and the genome size distribution for plant, respectively, both of which fit log-normal distribution (Fig 1A). Due to the additivity of normal distribution, the genome size distribution for animal and plant fit log-normal distribution (Fig 1A).

### Observation of genome size layout

For each taxon, I obtain the logarithmic mean of the genome sizes *E* and the standard deviation of the logarithmic genome sizes *σ*. The variation range of genome sizes can be estimated by the interval between *E* − *σ* and *E* + *σ*. Most studies in the literatures focus on the variation of genome sizes (Gregory 2005); however there was a lack of study on the relationship between *E* and *σ*. I introduce a two-dimensional layout plain to study the variation pattern of genome sizes for a taxon. The abscissa of the layout plain is the logarithmic mean of the genome sizes *E*, while the ordinate of the layout plain the standard deviation of the logarithmic genome sizes *σ*. I found that the standard deviation *σ* tends to decrease along the evolutionary direction. It is customary to set the upward direction of the ordinate as the evolutionary direction, so the standard deviation *σ* decreases along the upward direction of the ordinate in the layout plane. It is convenient to study the relationship between *E* and *σ* in the layout plain (Fig 1C, 1D).

The statistical features of genome sizes can be illustrated by the genome size layout. Genome size layout for a taxon is defined in the layout plain by plotting variation ranges of genome sizes for all the lower-ranked taxa with respect to the standard deviations of logarithmic genome sizes of species for the lower-ranked taxa respectively (Fig 1C, 1D). Based on the genome sizes and taxonomy in the Animal Genome Size Database and the genome sizes and taxonomy in the Plant DNA C-values Database (Gregory et al. 2007), the genome size layout for animal is obtained by plotting the variation ranges of the 19 animal taxa (Fig 1C). The genome size layout for angiosperm is obtained by plotting the variation ranges of the 53 angiosperm taxa (Fig 1D), where the 34 Dicotyledoneae taxa and the 19 Monocotyledoneae taxa are distinguished by different colour.

Two types of genome size layouts are defined as follows. In the case of V-shaped layout, the variation ranges are spread narrowly in lower layout plain (corresponding to large *σ*), while the variation ranges are spread widely in upper layout plain (corresponding to small *σ*). In the case of Λ-shaped layout, the variation ranges are spread widely in the lower layout plain, while the variation ranges are spread narrowly in the upper layout plain. I find that the genome size layout for animal is roughly Λ-shaped (Fig 1C) and the genome size layout for angiosperm is V-shaped, where most genome size ranges for Dicotyledoneae taxa slants to the left, while most genome size ranges for Monocotyledoneae taxa slants to the right (Fig 1D).

### Simulation of the log-normal distribution of genome sizes

Small- and large-scale duplications contribute much to the genome size evolution (Gregory 2005), whose contribution can be quantitatively estimated by a genome size growth rate. From a statistical perspective, the trend in genome size evolution follows an exponential growth trend. The change of genome size *δS* is proportional to the genome size *S* itself: *δS* = *Sλδt*, where *t* is time and λ is the genome size growth rate that is independent of genome size *S*. So the trend of genome size evolution obeys the exponential growth trend: *S*(*t*) = *S* exp(*λt*). The exponential growth trend in genome size evolution was suggested by studying the relationship between the origin time of taxa and the estimated genome sizes (Sharov 2007); and it was also obtained in a formula to estimate the genome sizes (Li and Zhang 2010). Here, I obtained the exponential growth trend in genome size evolution based on the relationship between the logarithmic mean of genome sizes of species in 7 taxa and the origin time of the 7 taxa respectively (Fig 1B), where the origin times for the 7 taxa are as follows respectively: diploblastica, 560 Million years ago (*Ma*); protostomia, 542 *Ma* (Ediacaran-Cambrian); deuterostomia, 525 *Ma;* bryophyte, 488.3 *Ma* (Cambrian-Ordovician); pteridophyte, 416.0 *Ma* (Silurian-Devonian); gymnosperm, 359.2 Ma(Devonian-Carboniferous); angiosperm, 145.5 *Ma* (Jurassic-Cretaceous) (Willis and McElwain 2014; Gibling and Davies 2012; Couvreur et al. 2011).

The genome size evolution can be regarded as a stochastic process. When considering the stochastic impact on genome size growth, I obtain *δS* = *Sλδt + Sλδn*, where *n* is a standard normal random variable. So the genome size evolution obeys the stochastic equation *S*(*t*) = *S* exp(*λt* + *λn*). I propose a log-normal stochastic process model (Ghanem 1999) to simulate both the observation of the lognormal distribution of genome sizes and the observation of the exponential growth rate of genome size evolution. In the model, the genome size *S* increases step by step at certain probability *P* from the initial value *S*(1) = 1 base pair at step 1 to the final value *S*(*N*) base pairs at step *N*, where *N* is the maximum number of steps. From step *n* − 1 to step *n* (*n* = 2, 3,…, *N*), the genome size doubles at probability *P*, or remains the same at probability 1 − *P*. The net result leads to a certain expected growth rate, which ensures that the genome size tends to increase exponentially. This is a stochastic process, where the probability *P* brings about the log-normal distribution of genome size *S*(*N*). The exponential growth rate in genome size evolution is constant in the simulation, when the probability *P* keeps constant. The whole process from step 1 to step *N* can be divided into *M* stages. I chose *M* steps *n* = *t* * (*N/M*)*, t* = 1, 2,…, *M* in the process, where *M* is the number of stages and *t* can be interpreted as the time in genome size evolution. Thus I obtained *M* genome size distributions *S*(*t* * (*N/M*)) at the *M* time *t* = 1, 2,…, *M* respectively. In the simulation, these genome size distributions *S*(*t* * (*N/M*)) are calculated based on the results of genome sizes at time *t* by running the process from 1 to *n*(*t*) for sufficiently many times. Due to the probability *P*, the values of genome sizes changes randomly in different runnings, but the obtained genome size distributions follow log-normal distribution. The logarithmic means of genome sizes for the simulated taxa increase with respect to time *t* due to the increasing numbers of steps *n*(*t*). These obtained *M* genome size distributions *S*(*t* * (*N/M*)) can be interpreted as the genome size distributions of one taxon at different stages (*t* = 1, 2,…,*M*) in the evolution (Fig 2A).

I obtained 10 genome size distributions for a simulated taxon respectively based on the log-normal stochastic process model (Fig 2A). The parameters in the simulation (Fig 2A) are as follows. The number of steps is set as *N* = 1, 000, 000, the number of time intervals is set as *M* = 10, and the probability is set as *P* = 0.2. 10 genome size distributions are calculated at time *t* = 1, 2, …, 10, respectively, by running from 1 to *t* * 1000, 000 for sufficiently many times. All the genome size distributions by the simulation fit log-normal distributions (Fig 2A). The exponential growth rates in genome size evolution are common in the simulation due to the same probability *P*. The logarithmic mean of genome sizes increases approximately linearly with respect to time *t* (Fig 2B), which means that the genome size increases exponentially. So the observation of the exponential trend in genome size evolution (Fig 1B) can be explained by the log-normal stochastic process model (Fig 2B). The simulation results in this paper explain the statistical observations of genome sizes qualitatively rather than quantitatively. The obtained logarithmic genome sizes can fit the observed logarithmic genome sizes of true species if adjusting the number of steps *N* properly in the simulations. So, it is convenient that the values of the abscissa are not shown explicitly so that the abscissa stands for the linear variation of logarithmic genome size qualitatively (Fig 2, 3).

**Figure 3:**
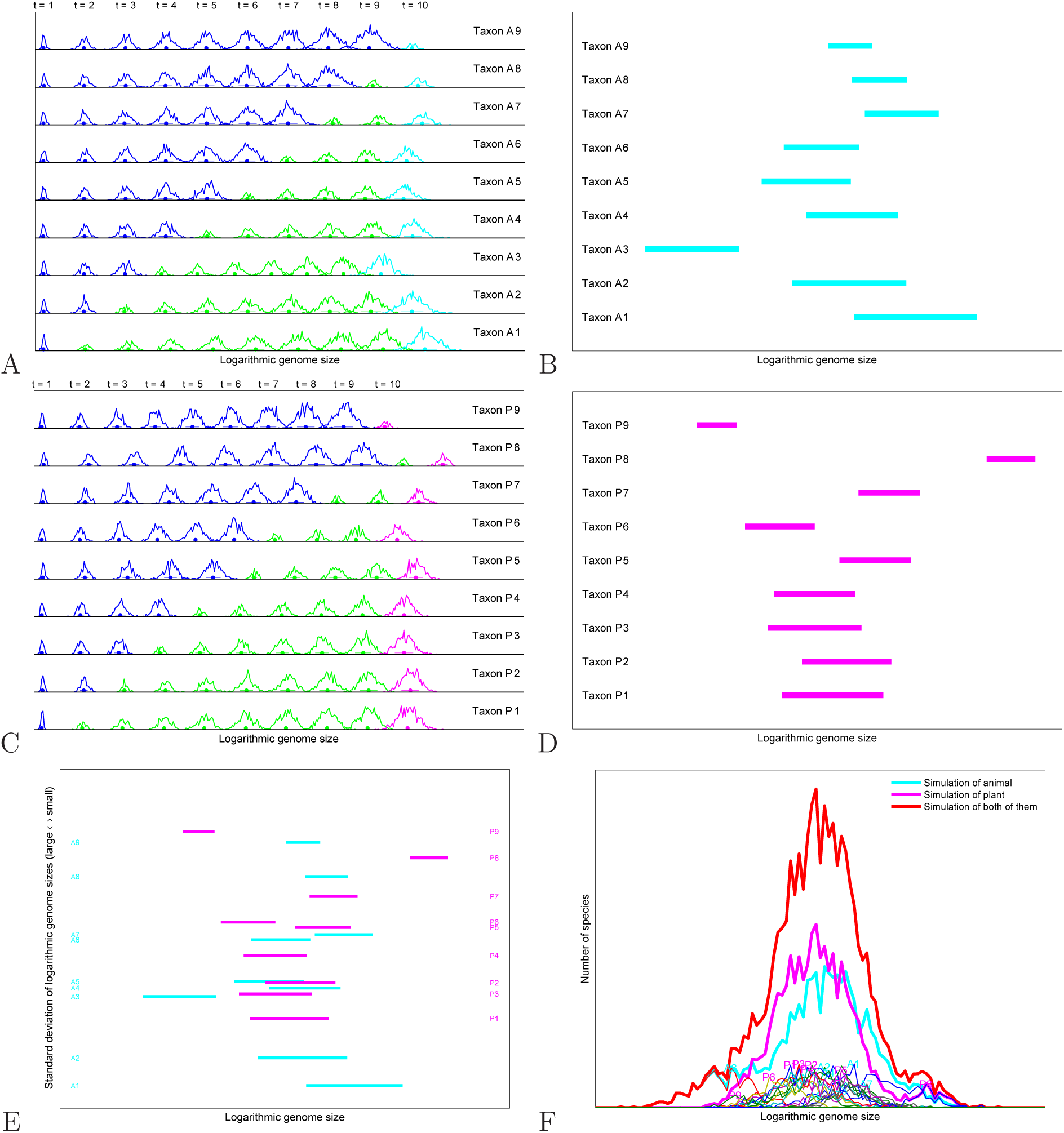
Explanation of the statistical features of genome sizes for animal and plant. (A~B) Explanation of the Λ-shaped genome size layout for animal. **A** Simulation of the genome size evolution for animal, where the probability for ancestor taxa is constant *P_a_* = 0.2, and the probability for descendant taxa *Pd* varies randomly from 0.175 to 0.225. **B** The variation ranges of genome sizes for the simulated animal taxa *Taxon A1* ~ *A*9, based on the results at time *t* = 10 in Fig 3A. (C~D) Explanation of the V-shaped genome size layout for angiosperms. **C** Simulation of the genome size evolution for angiosperms, where the probability for descendant taxa is constant *Pd* = 0.2, and the probability for ancestor taxa *P_a_* varies randomly from 0.175 to 0.225. **D** The variation ranges of genome sizes for the simulated taxa *Taxon P*1 ~ P9, based on the results at time *t* = 10 in Fig 3C. **E** The Λ-shaped genome size layout for animal and the V-shaped genome size layout for angiosperms based on the results in Fig 3B and 3D. **F** Simulation of the log-normal genome size distributions for animal, plant, and Eukarya, where the genome size distribution for animal is obtained by summing up the genome size distributions of *Taxon A*1 ~ *A*9 at time *t* = 10; the genome size distribution for plant is obtained by summing up the genome size distributions of *Taxon P*1 ~ *P*9 at time *t* = 10; and the genome size distribution for Eukarya is obtained by summing up the genome size distributions for animal and plant.

### Simulation of Λ-shaped and V-shaped genome size layouts

The statistic features of genome sizes in observation (Fig 1C, 1D) can be explained by developing the stochastic process model from single-taxon model to multi-taxa model. Essentially, the probabilities *P* are no longer constant in the multi-taxa model, which bring about different genome size growth rates. Hence the multi-taxa simulation can be achieved. The Λ-shaped and V-shaped genome size layouts in observation can be simulated if choosing different genome size growth rates for different stages in genome size evolution. The difference between the simulation of Λ-shaped layout and the simulation of V-shaped genome size layout results from the different assignment schemes for the probabilities *P* between the ancestor taxa and the descendant taxa.

The genome size layout is simulated based on the simulation of genome size distributions of many taxa by the mult-taxa model. There was only one taxon in the simulation in the previous subsection (Fig 2A). Now 9 taxa *Taxon k* (*k* = 1, 2,…, 9) are introduced in the present subsection according to the different assignment for the probability *P* (Fig 2C-2H). For each taxon *Taxon k*, the probability *P* is assigned differently between the stages from *t* = 1 to *t* = *k* (corresponding to ancestor taxa) and the stages from *t = k* + 1 to *t* =10 (corresponding to descendant taxa). The separation time between the ancestor taxa and descendant taxa for *Taxon k* (*k* = 1, 2,…, 9) are at time *t* = *k* + 1 (*k* = 1, 2,…, 9), respectively (Fig 2C). In this picture, only descendant taxa at stage *t* = 10 in the evolution are interpreted as the taxa in modern time (referred as modern taxa), while all the other taxa in the stages from *t* = 1 to *t* = 9 can be regarded as inaccessible from modern time. The genome size distributions for *Taxon k* (*k* = 1, 2,…, 9) are obtained based on the simulation results of modern taxa (denoted in red). All the taxa *Taxon k* (*k* = 1, 2,…, 9) at present *t* = 10 have a common ancestor taxon at time *t* = 1 (Fig 2C). The genome size layout for the corresponding higher-ranked taxon is obtained based on the 9 genome size distributions of the taxa *Taxon k* (*k* = 1, 2,…, 9).

In a special case when the probability is assigned constantly *P* = 0.2 for both ancestor taxa and descendant taxa, both the exponential growth rates for ancestor taxa and descendant taxa keep constant. Hence the logarithmic means of genome sizes for the 9 taxa *Taxon k* (*k* = 1, 2,…, 9) at time *t* =10 are constant. However, the standard deviations of the logarithmic genome sizes for the 9 taxa *Taxon k* decrease from *k* = 1 to *k* = 9, because the separation time changes from early to late for the taxa *Taxon k* from *k* = 1 to *k* = 9. Thus the genome size layout for the corresponding higher-ranked taxon can be obtained based on the variation ranges of genome sizes and the standard deviations of the logarithmic genome sizes for the 9 taxa *Taxon k*. The obtained genome size layout is of an isosceles triangle shape. The earlier the separation time between the ancestor taxon and descendant taxon is, the broader the variation range of genome size is. Roughly speaking, the earlier the taxa originated, the more the standard deviations of logarithmic genome sizes for taxa are. This result is helpful to understand the distributional pattern of genome sizes.

If the probability *P* is assigned differently for ancestor taxa and descendant taxa, the above isosceles triangle shaped genome size layout may change into scalene triangles. Thus the Λ-shaped and V-shaped genome size layouts in observation can be simulated (Fig 2E, 2H). In the case of Λ-shaped genome size layout (Fig 2E), the probability for the ancestor taxa is assigned by a constant *P_a_* = 0.2, while the probability for the descendant taxa *P_d_* is adjustable. When the probability *P_d_* is assigned by *P_d_* = 0.175, a right-slanted scalene triangle is obtained, which forms the left side of the Λ-shaped genome size layout (Fig 2C, 2E). When the probability *P_d_* is assigned by *P_d_* = 0.225, a left-slanted scalene triangle is obtained, which forms the right side of the Λ-shaped genome size layout (Fig 2D, 2E). Thus, the Λ-shaped genome size layout is obtained by simulation (Fig 2E). The simulation indicates a relationship between the genome size layout and the phylogeny of life. According to the simulation, the taxa *Taxon k* (*k* = 1, 2,…, 9) have common ancestor with same genome size growth rate. So the Λ-shaped genome size layout of a taxon indicates that the corresponding taxon is monophyletic.

In the case of V-shaped genome size layout (Fig 2H), the probability for the descendant taxa is assigned by a constant *P_d_* = 0.2, while the probability for the ancestor taxa *P_a_* is adjustable. When the probability *P_a_* is assigned by *P_a_* = 0.225, a left-slanted scalene triangle is obtained, which forms the left side of the V-shaped genome size layout (Fig 2F, 2H). When the probability *P_a_* is assigned by *P_a_* = 0.175, a right-slanted scalene triangle is obtained, which forms the right side of the V-shaped genome size layout (Fig 2G, 2H). Thus, the V-shaped genome size layout is obtained by simulation (Fig 2H). This simulation also indicates a relationship between the genome size layout and the phylogeny of life. According to the simulation, the taxa *Taxon k* (*k* = 1, 2,…, 9) have different ancestors with different genome size growth rates. So the V-shaped genome size layout of a taxon indicates that the corresponding taxon is polyphyletic.

## Results and discussion

### Explanation of the genome size layout for animal

The genome size layout for animal is about Λ-shaped in observation (Fig 1C). In the simulation of the genome size layout for animal (Fig 3A, 3B), the probability for ancestor taxa *P_a_* is still assigned by a constant *P_a_* = 0.2, while the probability for descendant taxa *P_d_*(*k*) (*k* = 1, 2,…, 9) is assigned by a random variable ranging between 0.175 and 0.225 (Fig 3A). 9 animal taxa *Taxon A*1 ~ *A*9 are simulated in total, where the genome size growth rates vary randomly with *P_d_*(*k*) (*k* = 1, 2,…, 9) (Fig 3A). Thus, the simulated genome size layout for animal is obtained (Fig 3B) based on the logarithmic means of genome sizes and standard deviations of logarithmic genome sizes of the 9 animal taxa *Taxon A*1 ~ *A*9 (Fig 3A). Although the simulated genome size layouts change randomly due to the randomly varying *P_d_*(*k*), the genome size layouts are always Λ-shaped due to the constant *P_a_*. The features of genome size layout for animal in observation (Fig 1C) are explained by the simulation (Fig 3B): both the genome size layout in observation (Fig 1C) and the genome size layout by simulation (Fig 3B) are Λ-shaped, where the variation ranges of genome sizes are large in early stages (lower in the genome size layout), while they become small in the late stages (upper in the genome size layout).

The simulation indicates that the animal kingdom is monophyletic, according to the relationship between the genome size layout and the phylogeny of life. This result agree with the common sense on metazoan origination. Almost all phyla of animal originated together at the Cambrian explosion (Conway-Morris 1993, Conway-Morris 2003, Valentine 2001, Shu 2008). It is generally believed that animal is monophyletic (Wainright et al. 1993, Cavalier-Smith et al. 1996). A Λ-shaped genome size layout for animal indicates a common ancestor for animals (Fig 3A, 3B), which support its monophyletic origin.

### Explanation of the genome size layout for angiosperm

The genome size layout for angiosperm is about V-shaped in observation (Fig 1D). In the simulation of the genome size layout for angiosperm (Fig 3C, 3D), the probability for ancestor taxa *P_a_*(*k*) (*k* = 1, 2,…, 9) is assigned by a random variable ranging between 0.175 and 0.225, while the probability for descendant taxa *P_d_* is assigned by a constant *P_d_* = 0.2 (Fig 3C). 9 plant taxa *Taxon P*1 ~ *P*9, corresponding to *P_a_*(*k*), are simulated in total, which can also be interpreted as angiosperm taxa (Fig 3C). Thus, the simulated genome size layout for angiosperm is obtained (Fig 3D) based on the logarithmic means of genome sizes and standard deviations of logarithmic genome sizes of the 9 angiosperm taxa *Taxon P*1 ~ *P*9 (Fig 3C). The simulated V-shaped genome size layouts also change randomly with the random variable *P_a_*(*k*). The features of genome size layout for angiosperm in observation (Fig 1D) are explained by the simulation (Fig 3D): both the genome size layout in observation (Fig 1D) and the genome size layout by simulation (Fig 3D) are V-shaped, where the variation ranges of genome sizes are small in early stages (lower in the genome size layout), while they become large in the late stages (upper in the genome size layout).

The separation between Dicotyledoneae taxa and Monocotyledoneae taxa in the genome size layout in observation (Fig 1D) can be explained by the simulation, due to the lack of a common ancestor for angiosperms. Most genome size ranges for Dicotyledoneae taxa slants to the left in the V-shaped genome size layout (Fig 1D), which indicates that the genome size growth rates for the ancestor of Dicotyledoneae taxa are large (Fig 2F, 2H). While most genome size ranges for Monocotyledoneae taxa slants to the right (Fig 1D), which can be simulated by the taxa whose ancestor taxa have small genome size growth rates (Fig 2G, 2H). The branching process between Dicotyledoneae taxa and Monocotyledoneae taxa results from the differences between their ancestors, according to the simulation of V-shaped genome size layout (Fig 2H).

There is a relationship between the type of the genome size layout and the phylogeny of life. The genome size layout for angiosperm is V-shaped, which differs from the Λ-shaped genome size layout for animal. Different evolutionary history for animal and angiosperm leads to their different genome size layouts. The simulation of genome size layout of angiosperm indicates that the angiosperm is polyphyletic. The genome size growth rates for the ancestors of the present angiosperm taxa are quite different, which results in the V-shaped genome size layout for angiosperm. The simulation result agrees with some reports in literatures. It is reported that the classes in angiosperms did not originate together (Willis and McElwain 2014, Bremer et al. 2009). The two angiosperm branches, the monocotyledons and dicotyledons, may originate independently. So the angiosperms would be polyphyletic (Stuessy 2004).

### Explanation of the genome size distributions for animal and plant

The stochastic process approach to understanding the genome size evolution can also explain the lognormally distributed genome sizes in observation. In the simulation of the genome size layout for animal, 9 genome size distributions for the taxa *Taxon A*1 ~ *A*9 have been obtained (Fig 3A). These genome size distributions roughly fit log-normal distributions, which obey the additivity rule of normal distribution. The genome size distribution for animal is obtained by summing up all the genome size distributions of the taxa *Taxon A*1 ~ *A*9. This simulated genome size distribution for animal also fits log-normal distribution (Fig 3F), which agrees with the observed genome size distribution for animal (Fig 1A).

Similarly, 9 genome size distributions for the taxa *Taxon P*1 ~ *P*9 have been obtained in the simulation of the genome size layout for plant (Fig 3C). These genome size distributions roughly fit log-normal distribution. The genome size distribution for plant is obtained by summing up all the genome size distributions of the taxa *Taxon P*1 ~ *P*9. This simulated genome size distribution for plant also fits log-normal distribution (Fig 3F), which agrees with the observed genome size distribution for plant (Fig 1A).

The simulated genome size distribution for Eukarya is obtained by summing up the simulated genome size distributions for animal and plant (Fig 3F), which agrees with the observed genome size distribution for Eukarya (Fig 1A).

## Acknowledgements

Supported by the Fundamental Research Funds for the Central Universities.

